# Small molecule inhibitor binds to NLRP3 and prevents inflammasome activation

**DOI:** 10.1101/2023.12.13.571573

**Authors:** Angela Lackner, Julia Elise Cabral, Yanfei Qiu, Haitian Zhou, Lemuel Leonidas, Minh Anh Pham, Alijah Macapagal, Sophia Lin, Emy Armanus, Reginald McNulty

## Abstract

Despite recent advances in the mechanism of oxidized DNA activating NLRP3, the molecular mechanism and consequence of oxidized DNA associating with NLRP3 remains unknown. Cytosolic NLRP3 binds oxidized DNA which has been released from the mitochondria, which subsequently triggers inflammasome activation. Human glycosylase (hOGG1) repairs oxidized DNA damage which inhibits inflammasome activation. The fold of NLRP3 pyrin domain contains amino acids and a protein fold similar to hOGG1. Amino acids that enable hOGG1 to bind and cleave oxidized DNA are conserved in NLRP3. We found NLRP3 could bind and cleave oxidized guanine within mitochondrial DNA. The binding of oxidized DNA to NLRP3 was prevented by small molecule drugs which also inhibit hOGG1. These same drugs also inhibited inflammasome activation. Elucidating this mechanism will enable design of drug memetics that treat inflammasome pathologies, illustrated herein by NLRP3 pyrin domain inhibitors which suppressed interleukin-1β (IL-1β) production in macrophages.

**One-Sentence Summary:** **NLRP3 cleaves oxidized DNA and small molecule drug binding inhibits inflammasome activation.**

## Introduction

Inflammation is the body’s temporary response to resolve an infection. Prolonged and uncorrected infection results in an overstimulated innate immune system that can lead to septic shock, organ failure, fibrosis, and cancer. The NOD-like receptor family pyrin domain containing 3 (NLRP3) inflammasome is a key mediator of tissue damage and pathogen(*1*) or toxicant(*2*) infection. The pathogenic role of NLRP3 was first discovered in humans harboring gain of function disease mutants(*3*). The NLRP3 inflammasome is a megadalton multi-protein complex consisting of NLRP3, ASC (apoptosis-associated speck-like protein containing a CARD (caspase activation and recruitment domain)) (*4*), pro-caspase-1, and NEK7 (Never in mitosis A (NIMA) – related kinase 7). Inflammasome autocleavage of pro-caspase-1 yields an active caspase-1 (p20/p10) heterotetramer. Caspase-1 protease activity leads to bioactive IL-1β, IL-18(*5*) and cleavage of gasdermin D (GSDMD) at Asp275 which causes GSDMD to oligomerize and insert itself into the membrane, where the electrostatic potential of its pore-lining residues allows selective transport of compatibly charged molecules which include IL-1β, IL-18, ATP, inflammatory caspases, cleaved GSDMD, and other cytokines and chemokines(*6*). These molecules amplify the immune response and recruit new immune cells that attempt to ameliorate the pathogen or danger signal. Extended efflux of molecules from macrophages causes a loss of cell membrane integrity, ultimately leading to cell death.

Canonical NLRP3 inflammasome activation consists of a 2-step activation process(*7*). Transcriptional priming can occur through bacterial LPS binding to TLR2/4 receptor, which leads to NF-κB induced expression of NLRP3 inflammasome subunits and pro-IL-1β(*8*). Reactive oxygen species (ROS) production in the mitochondria results in cytidine/uridine monophosphate kinase 2 (CMPK2) causing mitochondrial transcription(*9*). The unraveling of mtDNA required for transcription exposes the DNA to ROS which causes oxidation of mitochondrial DNA. Elimination of oxidized DNA by human 8-oxoguanine DNA glycosylase 1 (hOGG1) or mitophagy inhibits inflammasome activation. The presence of foreign and cytosolic signals including alum, asbestos, Sars-CoV2, or ATP overwhelm initial NLRP3 inhibition to promote inflammasome assembly and activation. The sustained presence of mtROS during mitochondrial replication further exacerbates oxidized mitochondrial DNA (Ox-mtDNA). Ox-mtDNA is cleaved to 500-650 bp fragments by flap endonuclease 1 (FEN1), which allows Ox-mtDNA to be exported to the cytoplasm via mitochondrial permeability transition pores (mPTP) and voltage-dependent anion channels (VDAC) where it can associate with NLRP3 and promote assembly and activation(*10*).

We have previously shown purified NLRP3 can directly bind Ox-mtDNA(*11*). Furthermore, NLRP3 pyrin shares a protein fold with hOGG1, which suggests NLRP3 might have both glycosylase and nuclease activity similar to hOGG1. This study aims to capture if NLRP3 has glycosylase activity and if preventing such activity with small molecules affects inflammasome activation. We show herein that NLRP3 can indeed cleave Ox-mtDNA. We also show small molecule drugs that prevent hOGG1 from interacting with oxidized DNA also prevent NLRP3 from interacting with Ox-mtDNA and block inflammasome activation in mouse macrophages.

## Results

### NLRP3 pyrin cleaves oxidized DNA

The active site of hOGG1 has been well characterized with the location of amino acids that interact with oxidized DNA illustrated by the crystal structure(*12*). Both sequence alignment and 3D superposition of hOGG1 and NLPR3 pyrin domain reveal a similar protein fold and many amino acids are identical or similar.

Glycosylases are the first enzymes in base excision repair (BER) and can recognize and excise oxidized bases from DNA lesions. All glycosylases that remove oxidized bases also cleave DNA by nucleophilic attack on the DNA backbone, generating a Schiff base protein-DNA intermediate via a lyase reaction using the amine group of a conserved lysine, thereby acting bifunctionally(*13*). This process results in cleavage of the N-glycosyl bond and release of the oxidized base, leaving an apurinic site (AP site) in the DNA which can be recognized by an AP endonuclease, APE1(*14*). APE1 creates a nick in the 5’ end of the AP site and adds a 3’ -OH which is required for DNA polymerase to add a new replacement base(*15*).

Since the fold and sequence of NLRP3 and hOGG1 are similar and NLRP3 pyrin can directly bind oxidized DNA (**Fig. 1A**)(*11*), we probed if NLRP3 pyrin domain could cleave oxidized DNA. NLRP3 pyrin incubated with 90-mer single stranded Ox-mtDNA produced a cleavage product which migrated lower than DNA alone (**Fig. 1B**). This data shows NLRP3 pyrin could behave like a bifunctional glycosylase; able to remove the oxidized base and cleave the backbone.

**Figure 1:**
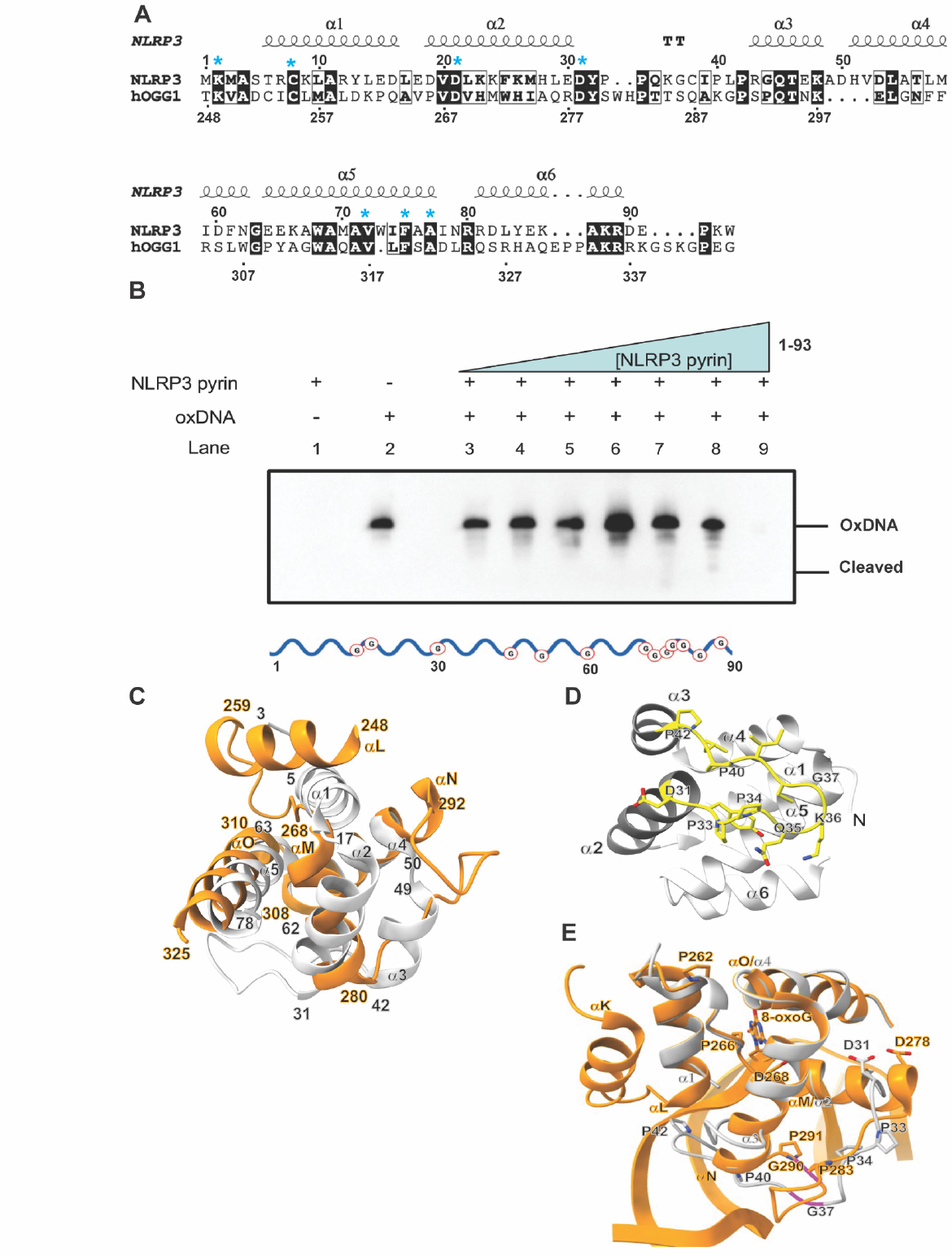
NLRP3 Pyrin domain shares a similar fold with hOGG1 and cleaves Ox-mtDNA. **(A)** NLRP3_(1-93)_ and hOGG1_(248-345)_ sequence alignment and secondary structure depiction. Cyan asterisks mark residues shared between NLRP3 and hOGG1 that play a pivotal role in binding and DNA base excision in hOGG1. **(B)** Biotinylated 90 base pair Ox-mtDNA incubated with increasing concentration of NLRP3 and probed with a streptavidin antibody shows DNA cleavage in a concentration dependent manner. **(C)** NLRP3 pyrin domain (grey) structure residues 3-78 with helices labelled α1-α5. Superposition of hOGG1 (orange) bound to oxidized DNA shown with corresponding helices αL-αO residues 248-325. **(D)** NLRP3 pyrin (PDBID: 7PZC, amino acids 1-91) contains a glycine/proline-rich region between amino acids 32 and 42, linking helices α2 and α3, and beginning with an aspartic acid (Asp31). NLRP3: grey, GPD-like loop: yellow **(E)** SWISS-MODEL projection of NLRP3 pyrin based on hOGG1 bound to Ox-DNA docked into the structure of hOGG1 bound to Ox-DNA (PDBID: 1EBM, amino acids 230-325). The GPD-like region of NLRP3 (amino acids 32-42) structurally aligns with a similar loop in hOGG1 (amino acids 279-291) which links helices αM and αN and also begins with an aspartic acid (Asp278). NLRP3: grey; hOGG1: orange.

To determine if the activity was specific to the pyrin domain of NLRP3, we performed the same assay with NLRP3 NACHT-LRR_(94-1034)_ construct, which lacks the pyrin domain. With increasing concentrations of protein incubated with 90-mer single stranded Ox-mtDNA cleavage was not seen, indicating that DNA cleavage is specific to the pyrin domain (**fig. S1**).

### NLRP3 and hOGG1 share protein fold features

Sequence comparison for NLRP3_(1-80)_ pyrin domain and hOGG1_(248-326)_ showed many residues that are either the same or similar between the two proteins. The sequence identity and similarity was 31.3% and 45.5%, respectively (**fig. S2**). Due to the moderate similarity in sequence, we compared the relative position of secondary structure in 3D (protein fold) of NLRP3 pyrin to hOGG1 (**Fig. 1C**). We first superposed NLRP3 pyrin_(1-81)_ and hOGG1_(248-326)_ (*16*).. To evaluate protein fold, the relative positions of NLRP3 pyrin helices α1-α5 was compared to corresponding positions of helices αL-αO for hOGG1 in the oxidized DNA-bound state (**Fig. 1C**) and the DNA-free state (**fig. S3**). We note NLRP3’s α3 is very short stretch from residues 42-49 which accounts for the extra helix topology. Helices α1 and αL did not align. The hOGG1 structure has a disordered stretch in the equivalent space of NLRP3 helix_α1_. The similarities between these two N-terminal locations include they are both predicted to be intrinsically disordered regions (IDR’s), which are regions predicted to convert between ordered and disordered (**fig. S4**)(*17*) . NLRP3_α2(17-31)_ and hOGG1_αM(268-280)_ traverse the same direction with different angles and have similar lengths of 19 Å and 17 Å, respectively. After the second helix, there is a loop for both proteins but in opposite directions. The NLRP3 loop (31-42) and hOGG1 loop (280-293) are similar in size, but the hOGG1 loop is slightly longer. Along the direction of the hOGG1 loop, NLRP3’s loop has a small helix, helix_α3(42-49)_, which has a strong kink (Ala49 to Asp50) causing a change in direction to continue along the same path as hOGG1_αN_. NLRP3_α3_ traverses the same direction as hOGG1 loop. Interestingly, both of the proteins have an IDR signature(*17*) for this region suggesting they are prone to convert between ordered and disordered secondary structures (**fig. S4**) according to the Database of Disordered Protein Predictions (D2P2)(*18*). The break in NLRP3_α3(42-49)_ directly connects to NLRP3_α4_, which aligns with hOGG1_αN_. Both proteins then turn to form NLRP3_α5(63-78)_ and hOGG1_αO(310-326)_ with those helices also aligned but slightly out of phase. This forms the end of hOGG1 C-terminus while NLRP3 pyrin domain has an additional helix, NLRP3_α6(80-90)_.

Human glycosylase OGG1 contains a Helix-hairpin-Helix (HhH) motif followed by a GPD region, a Gly/Pro-rich region which terminates with an Asp(*19*). The GPD loop in hOGG1 is critical for the cleavage of oxidized DNA(*12*). Changing the conserved Asp268 to Asn268 is a loss of function mutation for hOGG1(*20*). In our sequence alignment, NLRP3 Asp21 has identity to hOGG1Asp268 (**Fig. 1A**). However, NLRP3 Asp21 is not preceded by an HhH domain. Interestingly, NLRP3’s pyrin domain contains an HhH motif formed with a hairpin between helices α2-α3 (**Fig. 1D**). The 12-residue hairpin is GP-rich beginning with Asp31, Pro33, Pro34, Lys36 (apex), Gly37, Pro40, and Pro42. Unlike hOGG1 which has a GP-rich motif terminating with Asp268, NLRP3’s GP-rich hairpin begins with Asp31. Interestingly, this second GPD-like region that starts with Asp also maps to sequence alignment with hOGG1 residues 278-291 (**Fig. 1A**). In hOGG1, the equivalent region is about 180 degrees opposite the HhH motif. Similar to the HhH, this loop makes critical contacts with DNA which include Ala288(*21*) which, when mutated, can decrease substrate binding up to 60%(*22*). We generated a model of NLRP3 pyrin bound to oxidized DNA using SWISS-MODEL(*23*). The NLRP3 pyrin GPD-like region 31-42 traverses the same direction and orientation as hOGG1 278-291, in which both proteins start with Asp (**Fig. 1E**), confirming this region has propensity to interact with DNA.

### Glycosylase inhibitors prevent NLRP3 binding to oxidized DNA

It has been reported that NLRP3 can interact with oxidized DNA(*10, 11, 24*). The inhibitor TH5487 binds the 8-oxodG binding site of hOGG1 to prevent binding to oxidized DNA(*25*). Since the fold of NLRP3 and hOGG1 are similar, we probed if TH5487 could prevent NLRP3 pyrin from recognizing Ox-mtDNA. We found a concentration-dependent inhibition of NLRP3 binding to Ox-mtDNA in the presence of TH5487. Binding was inhibited with 100 μM TH5487 (**Fig. 2A-B**). No significant loss of binding to non-oxidized DNA was observed in the presence of the drug. We also tested the hOGG1 inhibitor SU0268(*26*) and saw a similar trend, in which the drug showed a concentration-dependent inhibition of NLRP3 binding to Ox-mtDNA, but did not effect binding to non-oxidized DNA (**Fig. 2C-D1**).

**Figure 2:**
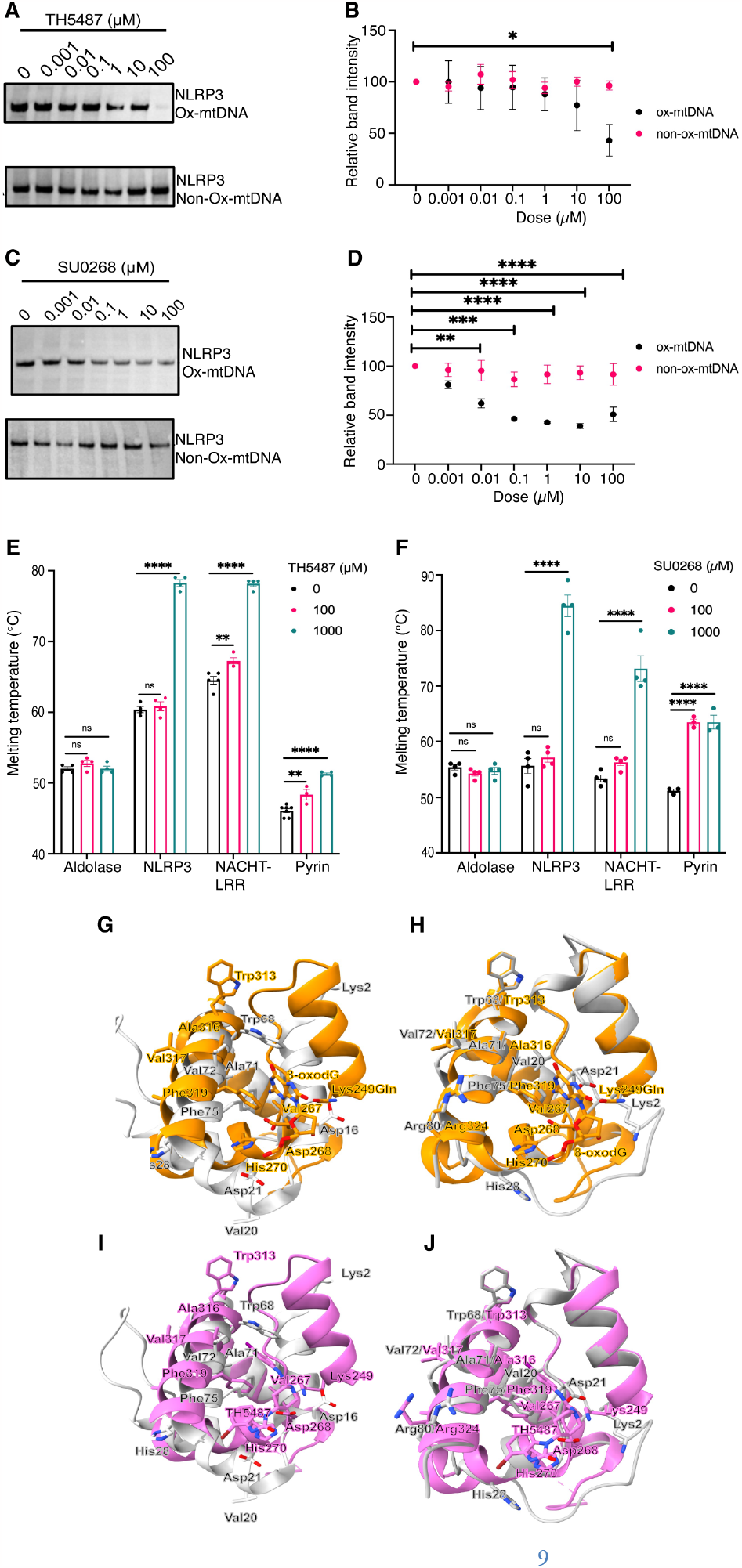
hOGG1 inhibitors bind NLRP3 and inhibit binding to Ox-mtDNA. **(A)** An anti-NLRP3 western blot on the bound fraction of NLRP3 to Ox- and non-Ox-mtDNA in the presence of TH5487 inhibitor from 0.001-100 µM. **(B)** Quantification of (A) shown as relative band intensity, N=4. **(C)** An anti-NLRP3 western blot on the bound fraction of NLRP3 to Ox- and non-Ox-mtDNA in the presence of SU0268 inhibitor from 0.001-100 µM.**(D)** Quantification of (C) shown as relative band intensity N=4, **P=0.0069. **(E)** Melting temperature data in presence of TH5487. N=4-8 NACHT-LRR **P= 0.0015. pyrin **P= 0.0036 **(F)** Melting temperature data in presence of SU0268. N=4 **(G)** NLRP3 pyrin (grey) PDBID: 7PZC docked into the structure of hOGG1 bound to Ox-DNA (orange) PDBID: 1EBM. **(H)** SWISS-MODEL generated NLRP3 pyrin bound to Ox-DNA (grey) docked into the structure of hOGG1 bound to Ox-DNA PDBID: 1EBM. **(I)** NLRP3 pyrin PDBID: 7PZC docked into the structure of hOGG1 bound to TH5487 (pink) PDBID: 6RLW. **(J)** SWISS-MODEL projection of NLRP3 pyrin based on hOGG1 bound to Ox-DNA docked into the structure of hOGG1 bound to TH5487 (PDBID: 6RLW). Error bars: mean +/-SEM. Analysis by one-way ANOVA, ****P <0.0001, ***P=0.0001, *P=0.211

### Glycosylase inhibitors directly bind NLRP3

To examine if TH5487 could directly bind NLRP3, we performed a thermal shift assay. The average melting temperature (T_m_) of NLRP3 shifted from an average value of 60.4 °C to 78.3 °C indicating the drug had directly bound to NLRP3 and delayed the T_m_ by an increase of 17.9 °C (**Fig. 2E**). We repeated this assay with the pyrin NLRP3_(1-93)_ and the NACHT-LRR NLRP3_(94-1034)_ domains. We found the pyrin domain could bind TH5487 as evidenced by the increase in T_m_ from 46.1 to 48.3 and 51.3 with 100 μM and 1 mM TH5487, respectively. Interestingly, the NACHT-LRR construct was also able to bind TH5487 and provide even more protein stability. The T_m_ was increased from 64.4 °C to 67.2 °C and 78.1 °C with 100 μM and 1 mM TH5487, respectively. Similar results were seen for drug SU0268 (**Fig. 2F**). As a control, we tested if the protein aldolase conferred similar changes to T_m_ in the context of TH5487. In these experiments, neither 100 μM nor 1 mM TH5487 caused a significant difference in the T_m_ of the protein compared to protein alone.

### NLRP3 and hOGG1 share active site residues

Upon superposition of NLRP3 pyrin_(1-81)_ with hOGG1_(248-326)_ many of the amino acids important in hOGG1 binding oxidized DNA (**Fig. 1C**) are also found in the active site for the 8-oxodG and the TH5487-bound states (**Fig. 2G-J and table S1**). The RMSD between NLRP3 pyrin and hOGG1 between 17 pruned atom pairs was 1.147 Å. Major differences appeared at the beginning of each model where the alpha carbon (Cα) for the critical Lys2 was 13.1 Å away from the corresponding residue Lys249 in hOGG1. Other amino acids in helices α3 – Leu54/299 and α4 -Trp 68/313, were separated by 0.6 and 3.8 Å, respectively. So, although many residues were in the same vicinity deviations were large enough to suggest NLRP3 pyrin would undergo a conformation change upon binding oxidized DNA. To generate a homology model of NLRP3 pyrin bound to oxidized DNA, we used the NLRP3_1-85_ sequence and hOGG1_248-326_ (PDBID: 1EBM) with the SWISS-MODEL software. A single model was generated with a MolProbity score of 1.78 and a Ramachandran favored percentage of 94.87% (**table S2**). The model illustrates a rearrangement of NLRP3 helix α1. The Lys2/249 distance decreased from 13.16 Å to 0.124 Å. Many residues including Ala4, Cys8, and Ala11 of NLRP3 all decreased to less than 1 Å apart in the NLRP3 model (**table S3**). All amino acids do not completely move to the same location. For example, Asp21/268 decreased from 7.551 Å to 6.322 Å apart. The corresponding to residues of NLRP3 in the active site of hOGG1 are Gln45/294, Trp68/313, Ala69/314, Ala71/316, Val72/317, Phe75/319, Ala77/321, Arg80/324, respectively. These residues all moved from an initial Cα-Cα distance of 2-20 Å to less than 1 Å apart.

### Blocking NLRP3 binding to oxidized DNA prevents inflammasome activation

It was previously demonstrated that targeting OGG1 to the mitochondria would decrease IL-1β production. Moreover, ablation of OGG1 leads to an increase of IL-1β since oxidized mtDNA has not been repaired. So, inhibiting OGG1 with TH5487, should cause an increase in IL-1β. Given we already observed that TH5487 could bind NLRP3, we examined what would happen to IL-1β in the presence of TH5487 under conditions that should promote inflammasome activation. Interestingly, we found that stimulation of NLRP3 inflammasome activation with LPS/ATP in the presence of TH5487 caused a decrease in IL-1β production with concentrations from 0.1-100 µM (**Fig. 3A-B**). Lower concentrations of TH5487 also decreased the amount of cleaved caspase-1 in the supernatant, and increased pro-caspase-1 in macrophages, which is consistent with a decrease in NLRP3 activation (**Fig. 3A and fig. S5**). The hOGG1 inhibitor, SU0268, showed the same significant decrease in NLRP3-dependent IL-1β dependent release, as well as a decrease in secreted caspase-1 (**Fig. 3C-D and fig. S6**).

**Figure 3:**
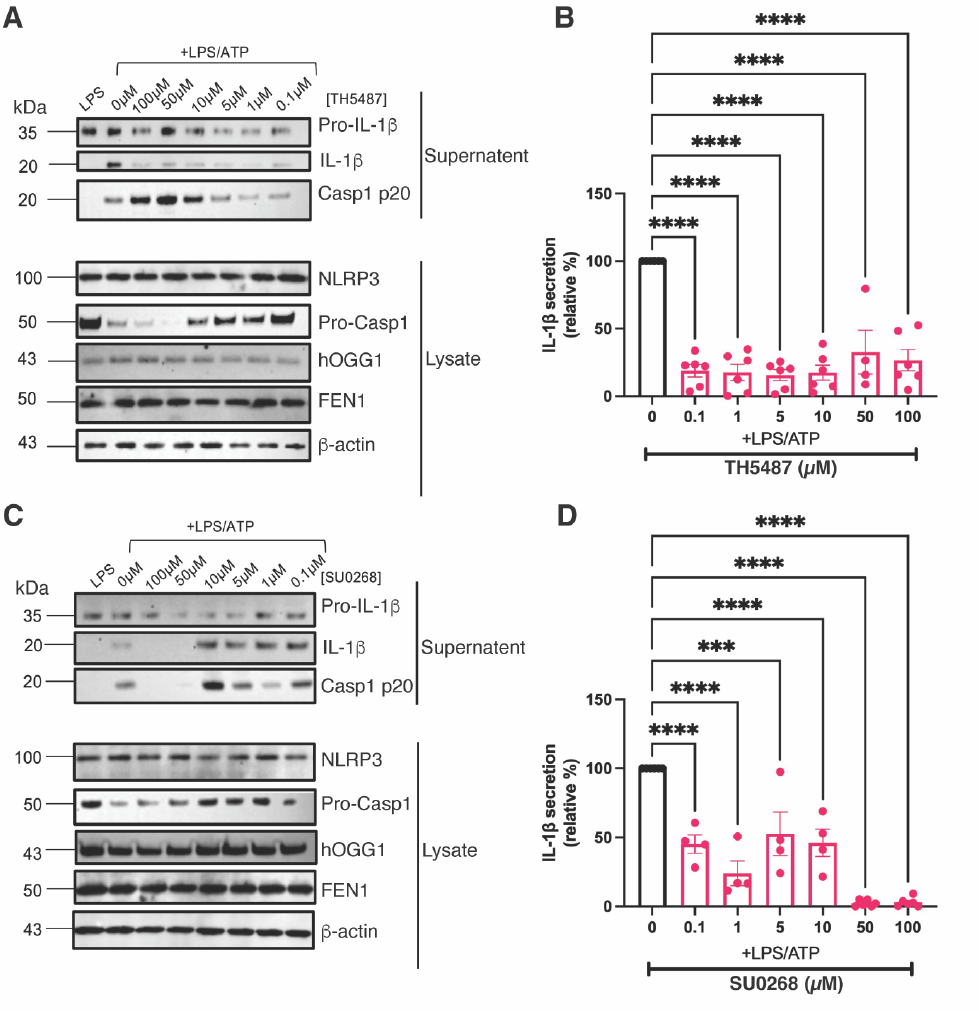
hOGG1 inhibitors TH5487 and SU0268 inhibit IL-1β secretion. **(A)** Representative western blots of inflammasome-related proteins in LPS/ATP treated or untreated immortalized mouse macrophages challenged with 0-100 µM TH5487. **(B)** Quantified relative amount of IL-1β in the supernatant of LPS/ATP stimulated immortalized mouse macrophages challenged with 0-100 µM TH5487. Error bars: mean +/-SEM, analyzed with one-way ANOVA. N=6, **** P<0.0001. **(C)** Representative western blots of inflammasome-related proteins in LPS/ATP treated or untreated immortalized mouse macrophages challenged with 0-100 µM SU0268 **(D)** Quantified relative amount of IL-1β in the supernatant of LPS/ATP stimulated immortalized mouse macrophages challenged with 0-100 µM SU0268. Error bars: mean +/-SEM, analyzed with one-way ANOVA. N=4, **** P<0.0001, *** P= 0.0002.

## Discussion

A connection between NLRP3 and Ox-mtDNA has been reported, however, the direct consequence and mechanism of NLRP3 binding Ox-mtDNA has not been established. Herein we describe that NLRP3 pyrin can cleave Ox-mtDNA. We describe the similarities in sequence, protein fold, and 3D structure with that of human glycosylase hOGG1, which functions in BER to remove oxidized guanine. The structure of hOGG1 and NLRP3 pyrin are similar enough such that small molecules which inhibit hOGG1 can also inhibit NLRP3 inflammasome activation in macrophages stimulated with LPS/ATP. Here, we present a model of NLRP3 bound to oxidized DNA and to two small molecule inhibitors, TH5487 and SU0268. Our model of NLRP3 cleaving oxidized DNA is consistent with major amino acids known to participate in DNA catalysis based on hOGG1 superposition and sequence alignment including Lys2 and Asp21, equivalent to hOGG1 Lys249 and Asp268, respectively. The ability of NLRP3 pyrin to not only excise the base and create an AP site, but cleave the N-glycosyl backbone suggests NLRP3 has bifunctional glycosylase activity.

Small molecules that inhibit the glycosylase’s ability to cleave oxidized DNA not only bind to NLRP3 (**Fig. 2A-F)**, but prevent NLRP3’s interaction with Ox-mtDNA and decrease NLRP3 inflammasome activation (**Fig. 3**). Since OGG1 ablation increases NLRP3 activation(*10*), use of these small molecule inhibitors to block NLRP3 activation may represent approaches to treat a variety of diseases for which NLRP3 inflammasome contributes, including non-alcoholic steatohepatitis (NASH), non-alcoholic fatty liver disease (NAFLD), hepatocellular carcinoma (HCC)(*27*), rheumatoid arthritis(*28*), keratitis(*29*) and over-active NLRP3 found in cryopyrin-associated periodic syndrome (CAPS) patients(*3*).

NLRP3 and hOGG1 have regions predicted to be intrinsically disordered, which by definition maintain the ability to convert between order and disorder depending on presence of binding partners. Using the sequence of the pyrin domain with AlphaFold2, which is documented to perform poorly with IDP’s, yields a structure that exactly matches the pyrin domain, since that structure is readily available in the Protein Data Bank(*30*). We found that SWISS-MODEL could produce a model that closely resembled hOGG1, which took into consideration not only sequence similarity, but order and disorder found in hOGG1 and the location of those residues in 3D. Superposition of NLRP3 pyrin with hOGG1 already shows several amino acids in NLRP3 that localize in the active site of hOGG1 in the proper vicinity to interact with oxidized DNA. The SWISS-MODEL generated structure of NLRP3, which represents the state of NLRP3 bound to oxidized DNA, not only has better location of amino acids in the active site, but the critical Lys2 need to form nucleophilic attack and cleave oxidized DNA more closely matches the hOGG1 structure. This change is enabled by a rearrangement of NLRP3_α1_ which has the propensity to stretch by unraveling α1-α2 transition (**Fig. 2G-J and fig. S4**).

It is quite interesting that the extramitochondrial presence of Ox-mtDNA, but not nuclear DNA, can induce arthritis when injected into mice(*31*). Mitochondrial DNA is normally well protected via TFAM packaging into nucleoids(*32*), however mtDNA is exposed during CMPK-dependent mitochondrial replication which allows easy oxidation of D-loop mtDNA. The nuclease FEN1 cleaves Ox-mtDNA into smaller fragments which can the exit the mitochondria via mPTP and VDAC channels(*10*). Once in the cytoplasm, the oxidized DNA has been demonstrated to bind to cyclic GMP-AMP synthase (cGAS) when present in large amounts(*33-35*). The cGAS-STING activation may synergize with the activation of NLRP3 inflammasome and production of IL-1β and IL-18. Cytosolic DNA will eventually be expelled from the cell via gasdermin D-mediated pyroptosis. The released Ox-mtDNA could then cause increased pro-inflammatory signaling as a DAMP by binding to the transmembrane protein TLR9 that leads to NF-κB signaling and IRF7-mediated type I IFN(*36*). The preference for TLR9 interacting with oxidized DNA over non-oxidized DNA establishes a more robust TNF and IL-8 response(*37*). The ability of NLRP3 to interact and remove oxidized guanine from oxidized DNA may serve to reduce inflammatory signaling similar to hOGG1. But clearly, situations that overwhelm the cell with cytosolic Ox-mtDNA cause inflammasome activation(*38*). Future work will involve probing the molecular determinants of NLRP3–Ox-DNA interaction as it directly relates to inflammasome assembly using advanced methods in cryo-electron microscopy(*39*).

## Acknowledgements

Funding: This work was supported by the National Institutes of Health NIAID grant K22AI139444 to R.M.

## Author contributions

A.L performed drug binding pulldowns, thermal shift assays, and inflammasome activation studies. J.E.C. performed clevage assays and sequence alignments. Y.Q., H.Z., L.L., M.P., A.M., S.L., and E.A. helped J.C. and A.L. with Bradfords and western blots. R.M. and A.L. made molecular models. R.M. designed experiments and supervised the project. A.L., J.E.C, and R.M. conceived of the study. R.M. wrote the manuscript. A.L., J.E.C, and R.M. edited the manuscript with input from all authors.

## Notes

### Competing Interest Statement

The authors have declared no competing interest.

