## Supplement for "Small molecule inhibitor binds to NLRP3 and prevents inflammasome activation"

### Materials and Methods

Purification and Expression of $NLRP3_{1-93}$

Site-directed mutagenesis (Agilent #210518) was performed on wild type NLRP3 to introduce a stop codon following the $93^{rd}$ amino acid, isolating the pyrin domain as previously explained(*1*). Briefly, $NLRP3_{1-93}$ was cloned in the mammalian expression vector pcDNA3.1HisB. The plasmid was then sequenced, grown at large scale, and purified with PureLink HiPure Expi Megaprep. The protein was expressed in 250 mL of media using the $Expi293_{TM}$ Expression System (Thermo). Enhancers were added 16 hours after transfection. Once the cells reached viability of <80% live cells, they were harvested by spinning at 1200 rpm in a swinging bucket rotor (JS-4.750, Beckman). After centrifugation, the dead cells were aspirated, and the pellet was resuspended in cold PBS to remove residual media. The protein was purified with half of the cell pellet on the AKTA Advant. Cells were resuspended in lysis buffer containing (50 mM Tris-HCL pH 7.4, 1X protease inhibitor (Pierce), 0.1% SDS, 10% glycerol, and 1% Triton X-100). The lysate was then sonicated for 42 s in intervals of 2 s on and 8 s off. Lysate was then clarified by spinning at 100,000 x g for 1 hour and subsequently passed through a 0.45 µm filter prior to affinity chromatography. The HisTrap FF crude 5 mL column was equilibrated in Buffer A (20 mM Tris pH 7.4, 200 mM NaCl, 10% glycerol) and eluted with Buffer B (20 mM Tris pH 7.4, 200 mM NaCl, 10% glycerol, 500 mM imidazole, 1 mM DTT, and 0.5% NP-40). The column was washed with Buffer A and eluted in a 5-step gradient of Buffer B (5%, 15%, 25%, 50%, 100%). Peak fractions were analyzed on a total protein NuPAGE 4-12% Bis-Tris run at 200 V for 30 minutes. Fractions were further analyzed with PVDF membrane western blots blocked in 2.5% BSA and probed with monoclonal pyrin targeting antibody (Adipogen) at a dilution of 1:10,000. Based on the result from the above gels, fractions were pooled and loaded onto a HiLoad 16/600 Superose 6 size exclusion column (20 mM Tris pH 7.4, 200 mM NaCl, 10% glycerol, 1 mM DTT, and 0.5% NP-40). No distinguishable peaks exist due to the presence of NP-40 in the buffer, thus all fractions were run on SDS page and western blots, and the $NLRP3_{1-93}$ sample was further analyzed via Native 4-16% gel (Figure S7). Protein concentrations were checked via Bradford (Biorad) to a final concentration of roughly 1 mg/mL.

Wild-Type NLRP3 and NACHT-LRR Expression and Purification

Wild-type NLRP3 was cloned into the mammalian expression vector pcDNA3.1HisB. The plasmid was expressed in DH5α cells (New England Biolabs) and purified using the PureLink HiPure Plasmid Maxiprep Kit (Thermo Fisher). The protein was expressed using the Expi293 Expression System (Thermo Fisher) per the manufacturer's instructions. Briefly, cells were grown in Expi293 expression media until they reached a concentration of $3\times10^6$ cells per milliliter and sustained viability of ≥95% live cells. At that time, 1 µg of expression vector was transfected per every 1 mL of cells with Expifectamine reagent. Once the cells reached viability of ≤80% live cells, they were harvested by spinning at 300 rpm for 5 minutes. The supernatant/dead cells were aspirated from the top, and the pellet was washed with cold PBS. The cells were pelleted again at 1200 rpm for 5 minutes and lysed with 50 mM Tris pH 7.4, 1 mM PMSF, 1 x protease and phosphatase inhibitor, 300 mM NaCl, 0.1% SDS, 10% glycerol, and 1% Triton X-100. The lysate was sonicated for 42 s in intervals of 2 s on, 8 s off, then clarified by spinning at 100,000 × g for 60 minutes. The clarified lysate was passed through a

0.45 µm filter and purified using a HisTrap FF crude 5 mL column. The column was pre-equilibrated with 20 mM Tris, 200 mM NaCl, 10% glycerol, 1 mM DTT, and 25 mM imidazole, pH 7.4. After the sample was loaded onto the column, it was washed with 10 column volumes (CV) of the wash buffer above. Then, using a four-step gradient from 25-100%, the protein was eluted using 20 mM Tris, 200 mM NaCl, 10% glycerol, 1 mM DTT, and 500 mM Imidazole, at pH 7.4. Peak fractions were pooled and loaded onto a HiLoad 16/600 Superose 6 pg size exclusion column. The column was run in a buffer containing 20 mM Tris, 200 mM NaCl, 10% glycerol, and 1 mM DTT, pH 7.4. Peak fractions were checked by SDS page and western blot. Samples were diluted with LDS sample loading buffer and reducing agent (Invitrogen), each at a final concentration of 1 x. The samples were boiled at 90°C and run on a NuPAGE™ 4 to 12%, Bis-Tris 1 mM 15-well mini-gels at 200 V for 30 minutes. For the western blot, samples were transferred to PVDF membranes, and blocked with 2.5% BSA. NLRP3 was probed with an anti-NLRP3 antibody (AdipoGen) at a 1:10,000 dilution in 2.5% BSA in TBST. NACHT-LRR$_{(94\text{-}1043)}$ was probed with a different anti-NLRP3 antibody (Cell Signaling) at a 1:1000 dilution in 2.5% BSA. Blots were incubated with an HRP-linked anti-mouse (Cell Signaling) or rabbit (Cell Signaling) secondary antibodies at 1:1000 dilutions and imaged using the iBright 1500 Imaging system. Once peak fractions were identified, they were pooled and concentrated using 100 kDa cut-off spin concentrators at cycles of 2000 × g for 5 minutes.

DNA Cleavage Assay

The DNA cleavage assay was performed with the purified pyrin domain and NACHT-LRR$_{94\text{-}1043)}$ constructs. NLRP3 pyrin$_{(1\text{-}93)}$ was dialyzed into buffer (150 mM NaCl 20 mM Tris, 0.05% NP-40, 1 mM DTT pH 7.4) with a 10 kDa MWCO (Thermo). The NACHT-LRR$_{94\text{-}1043)}$ construct remained in the size exclusion buffer for the DNA cleavage assay. Ox-mtDNA, sourced from IDT, was diluted in DEPC water to 2.7 ng/µL. DNA was incubated with purified protein for 72-96 hours at 4 °C prior to being analyzed on a 4-20% Tris-Glycine gel (Thermo) with a total reaction volume of 25 µL. Samples were boiled in 10% formamide and Orange Loading Dye containing SDS (NEB) for 2 minutes at 90 °C. The gel was loaded with Orange Loading Dye (NEB) and pre-run for 30 minutes at 100 V. The gel was run with 5 µL of reaction for 30 minutes at 100 V and transferred to an Immobilon-P 0.45 µm PVDF membrane (Millipore). After being transferred, the DNA was UV crosslinked for 15 minutes and blocked in 2.5% BSA and TBST (150 mM NaCl, 20 mM Tris, 0.1% Tween pH 7.4). The membrane was then probed with streptavidin HRP-linked antibody 1:2000 (BD Parmingen #554066) and washed 3 times in TBST for 10 minutes. Chemiluminescence (Thermo) was performed, and the DNA imaged with IBright and further analyzed in IBright software.

SWISS-MODEL generation of hOGG1-based NLRP3 active model

Using ChimeraX the published structure of hOGG1 bound to Ox-DNA was opened (PDBID: 1EBM). The amino acid sequence of hOGG1$_{(249\text{-}325)}$ that aligned with the NLRP3 pyrin$_{(1\text{-}91)}$ domain was saved as a .pdb file. The sequence was uploaded to SWISS-MODEL in their User Template Modeling input option as the template file(*2*). Then, NLRP3 amino acids 1-85 were loaded as the target file (amino acids 1-90 did not generate a model). The SWISS-MODEL projection produced one model of NLRP3 based on the template hOGG1 structure. This model was further analyzed in comparison to wild-type NLRP3 and various hOGG1 structures using ChimeraX(*3*).

Basic Concentration-Dependent NLRP3 Pulldown

Dynabeads M-280 Streptavidin (Thermo Fisher) were removed from the storage solution and washed three times with binding buffer (50 mM Tris, 100 mM NaCl, 2 mM $MgCl_2$, 12% glycerol, pH 7.4) using a DynaMag-2 magnet. Biotinylated Ox-mtDNA sourced from IDT was diluted 4:400 from the 100 µM stock solution and incubated with the beads overnight at 4 °C while rotating. The next day, the beads were separated on the magnet and washed three times with binding buffer. During the washes, NLRP3 was serial diluted from the 1.5 mg/mL stock by 10, 100, and 1000 in binding buffer. After the final wash, protein of varying concentrations was added to the beads in various wells and incubated overnight at 4 °C. The next day, the beads were washed three times with binding buffer. To evaluate the amount of NLRP3 bound to the beads, the beads were resuspended in LDS sample loading buffer and reducing agent (Invitrogen), each at a final concentration of 1X. The samples were boiled at 90 °C and run on NuPAGE™ 4 to 12%, Bis-Tris 1 mM 15-well mini-gels at 200 V for 30 minutes. Samples were transferred to PVDF membranes, blocked with 2.5% BSA, and probed with an anti-NLRP3 antibody (AdipoGen) at a 1:10,000 dilution in 2.5% BSA in TBST. Blots were incubated with an HRP-linked anti-mouse antibody (Cell Signaling) at a 1:1000 dilution and imaged using the iBright 1500 Imaging system. The intensity of the bands at a 1 minute exposure time was used to decide which concentration of protein to use in the inhibitor competition assay. Non-diluted protein (1:1) was chosen as it gave the most visible band without overexposure at 1 minute.

Inhibitor Competition Pulldown

Dynabeads M-280 Streptavidin (Thermo Fisher) were removed from the storage solution and washed three times with binding buffer (50 mM Tris, 100 mM NaCl, 2 mM MgCl2, 12% glycerol, and pH 7.4) using a DynaMag-2 magnet. Biotinylated Ox-mtDNA sourced from IDT was diluted 4:400 from the 100 µM stock solution and incubated with the beads overnight at 4 °C while rotating. The following day, inhibitors TH5487 (Selleck Chemicals) and SU0268 (MedChemExpress) were serial diluted with binding buffer such that the addition of inhibitor at various concentrations was always 10% of the final volume. Then the inhibitor was incubated with the protein (NLRP3, NACHT-LRR$_{(94\text{-}1034)}$, or pyrin$_{(1\text{-}93)}$) at various concentrations from 1 nM to 100 µM at 4 °C for 1 hour. A control of protein with binding buffer was also incubated at 4 °C for 1 hour. During these incubations, the beads were separated on the magnet and washed three times with binding buffer to remove any unbound DNA. Once the inhibitor incubations were complete, each incubation was added to a well with beads in triplicate, mixed by pipetting up and down, and incubated at 4 °C overnight. The following day, the beads were separated on the magnet, the supernatant was removed, and the beads were washed three times with binding buffer. To evaluate the amount of NLRP3 bound to the beads, the beads were resuspended in LDS sample loading buffer and reducing agent (Invitrogen), each at a final concentration of 1X. The samples were boiled at 90 °C and run on NuPAGE™ 4 to 12%, Bis-Tris 1 mM 15-well mini-gels at 200 V for 30 minutes. Samples were transferred to PVDF membranes, blocked with 2.5% BSA. Pyrin$_{(1\text{-}93)}$ and NLRP3 westerns were probed with an anti-NLRP3 antibody (AdipoGen) at a 1:10,000 dilution in 2.5% BSA in TBST. NACHT-LRR$_{(94\text{-}1034)}$ westerns were probed with a different anti-NLRP3 antibody (Cell Signaling) at a 1:1000 dilution in 2.5% BSA. Blots were incubated with an HRP-linked anti-mouse (Cell Signaling) or anti-rabbit (Cell Signaling) secondary antibodies for 1 hour at a 1:1000 dilutions and imaged using the iBright 1500 Imaging system. The intensities of the bands were quantified using the iBright Analysis Software. The intensity values were plotted and analyzed using GraphPad Prism and a one-way ANOVA.

Protein Thermostability Assay

NLRP3, NACHT-LRR$_{(94\text{-}1034)}$, pyrin(1-93), and aldolase proteins were diluted to 1 mg/mL and challenged with 100 or 1000 µM TH5487 (Selleck Chemicals) or SU0268 (MedChemExpress) diluted in 20 mM Tris, 200 mM NaCl, 10% glycerol, and 1 mM DTT, pH 7.4 for 1 hour at 4 °C. Each protein was also incubated with buffer alone as a control. During this incubation, the assay plate was set up. Plates (BioRad, #HSP9601) were set up in the dark on ice using the Protein Thermal Shift Dye Kit (Thermo Fisher, #4461146) per the manufacturer's instructions. Briefly, protein thermal shift dye was diluted to 8X, and 2.5 µL was added to each sample well along with 5 µL of protein thermal shift buffer. Once the protein incubations were completed, 12.5 µL was added to the sample wells in quadruplicate. Wells were mixed by pipetting up and down 3 times with a multichannel pipette, then spun down at 1000 rpm for 2 minutes. The plate was kept in the dark and on ice until analysis. Melting temperature analysis was done using a CFX Duet Real-Time PCR system (BioRad, #12016265) and the CFX Maestro Software 2.3 (BioRad, #12013758). To set-up a thermal shift assay, all other steps of the standard RT-PCR run were removed, and a melt curve was interested. An initial 30 seconds at 4 °C cycle was inserted before the ramp step. The protocol was set to ramp at a ramp rate of 0.5 °C every 10 seconds from 4-95 °C. The FRET channel was assigned to each sample well, and the plate was inserted into the machine and run. The resulting melting temperatures were plotted and analyzed using GraphPad Prism and a one-way ANOVA.

Mouse Macrophage Cell Culture

Immortalized wild-type mouse macrophages were generously provided to us by Michael Karin at the University of California, San Diego. A frozen vial of cells at $1*10^7$ was thawed in a water bath and then resuspended in 50 mL of culture media (DMEM (Thermo Fisher) supplement with 10% heat-inactivated FBS (Sigma) and 1% Penicillin-Streptomycin (Thermo Fisher)). The cells were spun down at 300 x g for 5 minutes at 4 °C, the media was aspirated off the pellet, and the pellet was resuspended in 5 mL of culture media. The cells were counted by diluting 10 µL of cells with 10 µL of Trypan Blue Stain (Thermo Fisher), loading that dilution onto a Countess™ Cell Counting Chamber Slide (Thermo Fisher), and evaluated using a Countess™ 3 FL Automated Cell Counter (Thermo Fisher). Cells were then further diluted in culture media such that the final concentration was $0.2*10^6$/mL and plated onto 175 $cm^2$ culture flasks (Corning) where they were grown at 37 °C and 5 % humidity. Cells typically doubled in 24-48 hours, where they were then scraped from the bottom of the plate using Bio-One Cell Scrapers (Fisher Scientific) and then spun down, counted, and expanded as previously described.

Macrophage Inflammasome Activation with Inhibitors

The viability and concentration of a culture of immortalized mouse macrophages were checked as described above. Cells at viability >95% were diluted to a concentration of $0.5*10^6$ cells/mL, such that they would be at a concentration of $1*10^6$ cells/mL by the next day. Cells were split into 6-well TC-treated plates (Corning) with 2 mL of cells per well and allowed to adhere and grow overnight at 37 °C and 5 % humidity. The next day, 2 μL/mL of 500X lipopolysaccharide (Thermo Fisher) was added to each well for 1 hour. Next, LPS-only wells were harvested, and inhibitors TH5487 (Selleck Chemicals) and SU0268 (MedChemExpress) were serial diluted such that the addition of any concentration of inhibitor was 1% of the final volume of cells. Then the inhibitors were added at concentrations ranging from 0.1-100 μM for 1 hour. Next, 4 μM ATP was added to each well for 1 hour. Cells and supernatant fractions could then be isolated for viability and western blot analysis. To collect samples for western blot analysis, the supernatant was removed from each well and clarified by spinning at 3000 x g for 5 minutes using 0.22 μm spin filters (Corning). The cells left on the plate were washed with ice-cold PBS and then lysed with RIPA buffer (Boston BioProducts) supplemented with an EDTA-free protease/phosphatase inhibitor cocktail (Roche). The lysis took place for 5 minutes rocking at 4°C. The lysed cells were then collected into 1.5 mL tubes and spun at 14,000 x g for 15 minutes. 200 μL of the clarified lysate was removed and saved for western blot analysis. To evaluate proteins secreted from the cells (Caspase-1 p20 and IL-1β), the supernatant fractions were run on western blots. To evaluate proteins expressed inside the cell (NLRP3, pro-Caspase-1, FEN1, hOGG1, and β-actin), the lysate fraction was run. Samples were diluted with LDS sample loading buffer and reducing agent (Invitrogen), each at a final concentration of 1X. The samples were boiled at 90°C and run on NuPAGE™ 4 to 12%, Bis-Tris 1 mM 15-well mini-gels at 200 V for 30 minutes. Samples were transferred to PVDF membranes, blocked with 2.5% BSA in TBST, and probed with a primary antibody against the specific protein diluted to the manufacturer's recommendation in 2.5% BSA in TBST (**table S4**). Blots were incubated with an HRP-linked secondary antibody (either mouse or rabbit, depending on the species of the primary) and imaged using the iBright 1500 Imaging system. The intensities of the bands were quantified using the iBright Analysis Software. The intensity values were plotted and analyzed using GraphPad Prism and a one-way ANOVA.

Measurement of Macrophage Cell Viability After Treatment

After completion of the ATP incubation of the macrophage inflammasome activation assay described above, sample wells for viability readings were analyzed. The supernatant from these wells was aspirated and saved for western blots as described above. 1 mL of fresh culture media (DMEM (Thermo Fisher) supplement with 10% heat-inactivated FBS (Sigma) and 1% Penicillin-Streptomycin (Thermo Fisher)) pre-warmed to 37 °C was added to each well. The cells were scraped off the plates using new Bio-One Cell Scrapers (Fisher Scientific) for each well. The now resuspended cells were then moved into 1.5 mL tubes The cells were counted by diluting 10 μL of cells with 10 μL of Trypan Blue Stain (Thermo Fisher), loading that dilution onto a Countess™ Cell Counting Chamber Slide (Thermo Fisher), and evaluated using a Countess™ 3 FL Automated Cell Counter (Thermo Fisher). The percent viabilities were recorded in triplicate, and the values were plotted and analyzed using GraphPad Prism and a one-way ANOVA. The remaining cells were spun down at 300 x g for 5 minutes at 4 °C. The supernatant was aspirated, and the cells were washed with ice-cold PBS. To remove the PBS, the cells were spun again at 300 x g for 5 minutes at 4 °C and the PBS was aspirated. The cells were lysed with RIPA buffer (Boston BioProducts) supplemented with an EDTA-free protease/phosphatase inhibitor cocktail (Roche) for 5 minutes rotating at 4 °C. The lysate was clarified by spinning at 14,000 x g for 15 minutes. 200 µL of the clarified lysate was removed and saved for western blot analysis as described above.

**Supplementary Text**

NLRP3 Cleavage of Ox-mtDNA is specific to the pyrin domain

Due to the sequential and structural homologies between hOGG1 and NLRP3 pyrin domain, we wanted to explore the possible functional similarity between the two. We first determined that the pyrin domain of $NLRP3_{(1-93)}$ can cleave Ox-mtDNA. Next, we wanted to probe the catalytic specificity of this region. Using a construct lacking the pyrin domain ($NACHT\text{-}LRR_{(94-1034)}$), we identified that under native **(fig. S1A)** and denaturing **(fig. S1B)** conditions, the protein binds but does not cleave Ox-mtDNA. This suggests that there may be multiple sites in NLRP3 that can bind Ox-DNA, as we previously reported, but that the catalytic activity like hOGG1 is specific to the pyrin domain.

NLRP3 is sequentially and structurally similar to hOGG1

As explained previously, $NLRP3_{1-80}$ and $hOGG1_{246-326}$ have many identical and similar residues, 31.3% and 45.4% respectively **(fig. S2)**. Much of this similarity is comprised of conserved residues crucial in Ox-DNA binding and base excision in hOGG1. With this high sequence similarity, we examined the structural similarities of both proteins. We performed a superposition of NLRP3 $pyrin_{3-78}$ with $hOGG1_{248-325}$ not bound to Ox-DNA **(fig. S3)**. These two seemingly different proteins appear to have several striking similarities in helix orientation and length. We note that NLRP3's $\alpha_1$ and hOGG1 $\alpha_L$ do not align, however, programs PV2, PrDOS, and VSL2b predict disorder in NLRP3's helix α1 and Espritz-N and PV2 predict disorder in this region for hOGG1, suggesting that these regions are both able to convert between order and disorder upon binding substrate **(fig. S4)**. Each loop is further analyzed in the main text. Briefly, NLRP3's $\alpha_2$ and hOGG1's $\alpha_M$ travel in the same direction as $\alpha_2$ with NLRP3's 2 Å longer. NLRP3's $\alpha_3$ traverses the same direction as the hOGG1 $loop_{280-293}$, and both contain an IDR (**fig. S4**). NLRP3 $\alpha_3$ and $\alpha_4$ both align to hOGG1's $\alpha_N$. Lastly, $\alpha_5$ and $\alpha_O$ travel in the same direction with a similar length but are slightly out of phase.

To further compare the similarity of hOGG1 and NLRP3 $pyrin_{(1-93)}$, we generated a model of NLRP3 pyrin bound to Ox-DNA using SWISS-MODEL. This model suggests that NLRP3 rearranges $\alpha_1$ and stretches the α1- α2 transition. PV2 and VSL2b suggest this region is disordered **(fig. S4)** and thus can convert between disorder and order, supporting the SWISS-MODEL results.

As we explained in the main text, glycosylases have a Helix-hairpin-Helix (HhH) motif followed by a GP-rich region terminating with an invariant aspartic acid. As we discussed, hOGG1's terminating Asp is residue 268, aligning with NLRP3's Asp21. However, we report that NLRP3 contains a HhH-like motif with the hairpin beginning with the invariant Asp31. We further explain that hOGG1 contains a second GPD-like region comprised of residues 278-291. Espritz-

N, IU-Pred-S, and PV2 predict that this region in between α2-α3 of NLRP3 is intrinsically disordered. Similarly, Espritz-N, Espritz-X, IUPred-L, IUPred-S, PV2, PrDOS, VSL2b, and VLXT predict the region between the αM-αN of hOGG1 as a region of disorder **(fig. S4)**. This similarity in disorder suggests that this region may be able to bind Ox-DNA in NLRP3.

hOGG1 inhibitors specifically inhibit NLRP3 activation by inhibiting Ox-mtDNA binding

We have already shown that NLRP3 activation in mouse macrophages is inhibited by treatment with both hOGG1 inhibitors, TH5487 and SU0268 as evidenced by a decrease in IL-1β secretion (**Fig. 7 and Fig. 8**). However, we further probe the expression of other proteins involved in the NLRP3 inflammasome to define the inhibitory pathway. The TH5487 inhibitor had no effect on NLRP3, hOGG1, or FEN1 expression (**fig. S5A, D, E**) suggesting the inhibitor acts to directly inhibit NLRP3 activation. Treatment with this drug did show an increase in pro-caspase-1 in the cells, which suggests that pro-caspase-1 was not cleaved into active caspase-1, indicative of NLRP3 activation, and explaining the decrease in caspase-1 secretion (**fig. S5B, C**). We also test the viability of the cells to ensure the results are not due to a loss in cell viability. There is a loss in cell viability at the higher TH5487 concentrations (50 and 100 μM) suggesting the data at those doses should not be interpreted as specific to the TH5487 inhibitor (**fig. S5F**). However, we do see compelling data, at the lower dose ranges of TH5487, which we may attribute to effects caused directly by the drug, and not by a loss in cell viability. We see similar data with the inhibitor SU0268. As with TH5487 treatment, NLRP3, hOGG1, and FEN1 expression remained unchanged upon challenge with SU0268 **(fig. S6A, D, E**), while caspase-1 secretion is decreased, and internal pro-caspase-1 is increased compared to no-drug controls (**fig. S6B, C**). Interestingly, this drug did not elicit the same loss in cell viability at higher concentrations as seen with TH5487 (**fig. S6F**) suggesting results seen across the whole drug dose range tested may be attributed to the drug treatment.

Model of NLRP3 in putative active confirmation confers improved structural homologies to hOGG1 bound of Ox-DNA

There are clear structural similarities between the pyrin domain of NLRP3 and hOGG1. Interestingly, the homologies are similar between NLRP3 and both hOGG1 bound to Ox-DNA and hOGG1 bound to the inhibitor TH5487 (**table S1**). Though these structures are similar, they are not exactly homologous, as evidenced by their RMSD values of 1.132 Å and 1.147 Å respectively. This data led us to model NLRP3 in a putative active confirmation based on the structure of hOGG1 bound of Ox-DNA. Using SWISS-MODEL, we were able to successfully generate a model of NLRP3 pyrin$_{(1\text{-}85)}$ that attempted to map to the structure of the hOGG1 active site (amino acids 249-325) but also considered the steric constraints NLRP3's amino acids (**table S2**). Upon structural analysis, the model provided improved structural homologies to hOGG1 bound to both Ox-DNA and TH5487, with RMSD values of 0.256 Å and 0.358 Å, respectively (**table S3**). These new homology scores show the structural rearrangement NLRP3 might undergo to confer an active confirmation, as well as the accessibility this confirmation might have for both the inflammasome activator, Ox-mtDNA, and the putative inhibitor, TH5487.

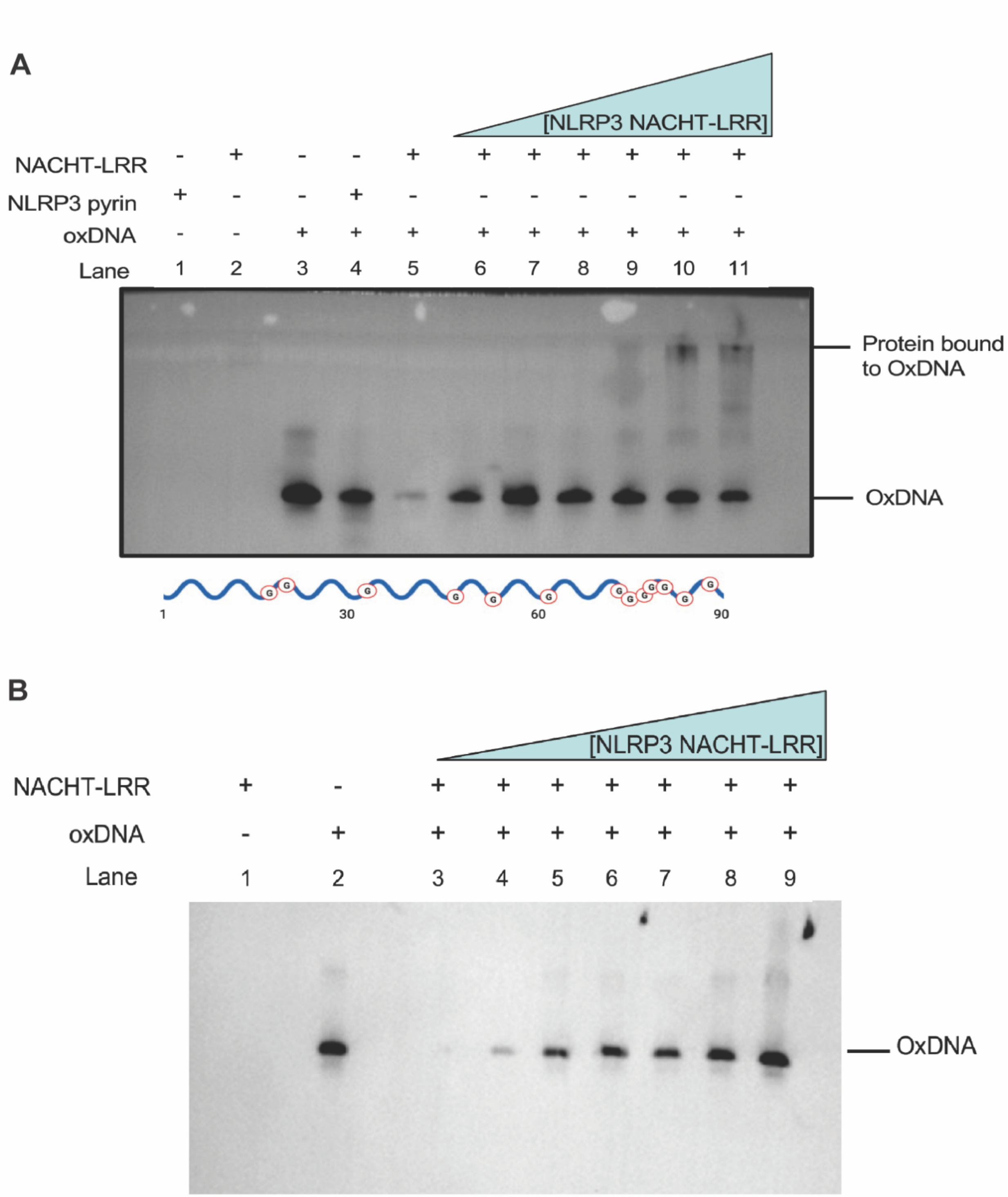


**figure S1: NLRP3 NACHT-LRR does not cleave oxidized DNA. (A)** The DNA cutting assay was performed on the NLRP3 NACHT-LRR$_{(94\text{-}1034)}$ under native conditions. The protein binds but does not cut Ox-mtDNA. **(B)** Experiment above was repeated under denaturing conditions

```
Matrix: EBLOSUM62
Gap penalty: 2.0
Extend penalty: 2.0
Score: 105.0
Sequence 1 length:80
Sequence 2 length:80
Alignment length: 99
Identity:      31/99 (31.31%)
Similarity:    45/99 (45.45%)
Gaps:          38/99 (38.38%)

1     GTKVADCICLMA--L-DKPQAVPVDV---HMWHIAQRDYSWHPTTSQAKG--PSP--QTN     50
        | |   |  |  | |      ||     | |    ||   |   | ||  | |  ||
1     -MKMASTRCKLARYLEDL-E--DVDLKKFKM-HL-E-DY---P--PQ-KGCIPLPRGQTE     47

51    K-E-LGNFFRSL---W-G-PYAGWAQAV-LFSADL-RQS                          80
      |          |     |   | || ||  | |   |
48    KADHV-D-LATLMIDFNGEEKA-WAMAVWIFAA-INR--                          80
```

**figure S2: hOGG1 and NLRP3 share many residues.** NLRP3 (1-80) and hOGG1 (246-326) have a sequence identity of 31.31% and a sequence similarity of 45.45% (VectorBuilder).

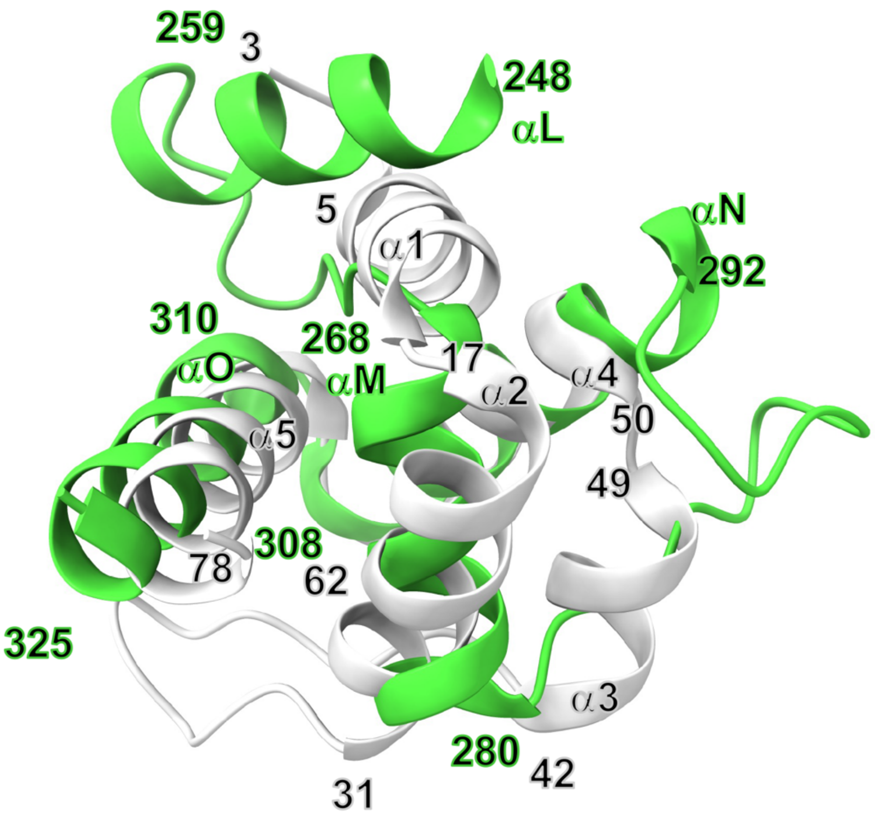


**figure S3: NLRP3 pyrin has similar fold to hOGG1 DNA-free structure.** NLRP3 pyrin domain$_{(3\text{-}78)}$ (grey) with helices labelled α1-α5. Superposition of hOGG1$_{(248\text{-}325)}$ in the absence of DNA (green) shown with corresponding helices αL-αO.

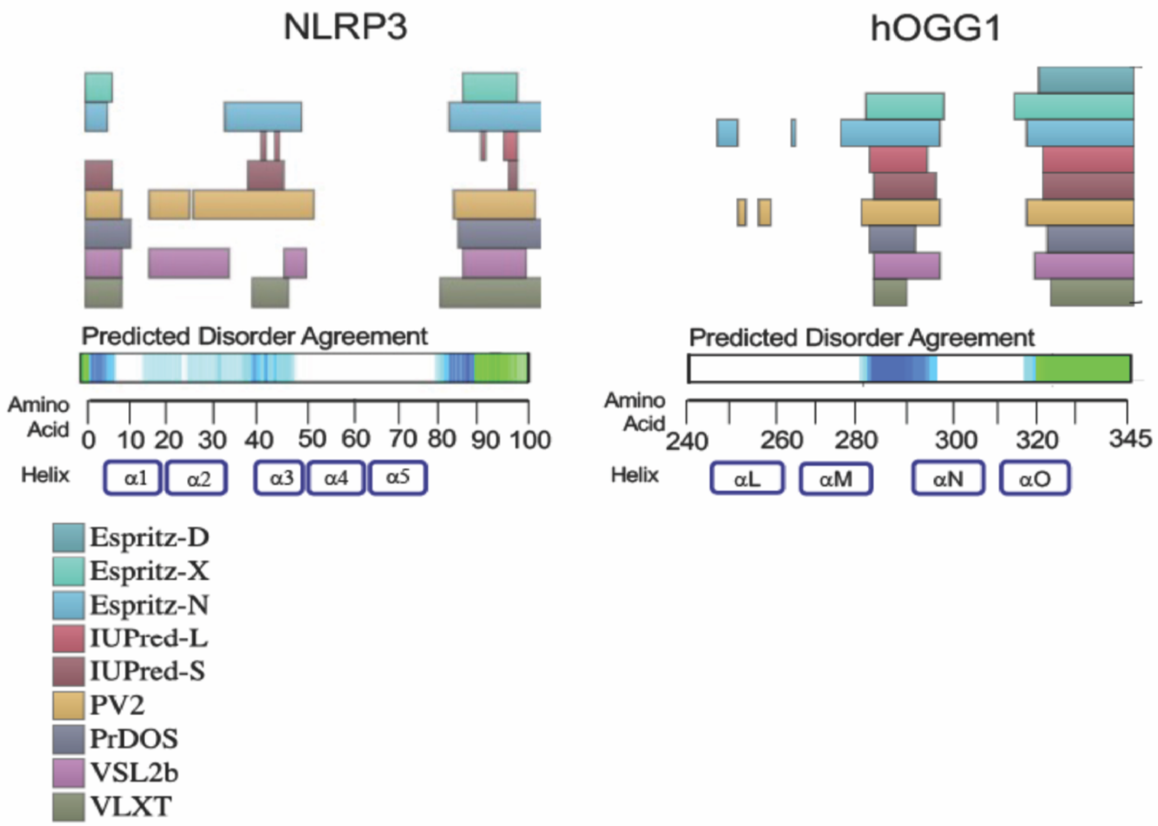


**figure S4: NLRP3 Pyrin and hOGG1 share similar regions of disorder.** Predicted disorder across several programs (color coded squares) in NLRP3 (left) and hOGG1 (right) corresponding to their amino acid sequence. Helix topology of NLRP3 is labeled α1-α5, and hOGG1 as αL-αM. Programs PV2, Espritz-N predict intrinsic disorder between NLRP3 helices α1-α2, and α2-α3 which matches the model of NLRP3 pyrin in the DNA bound state **(Fig. 2)**.

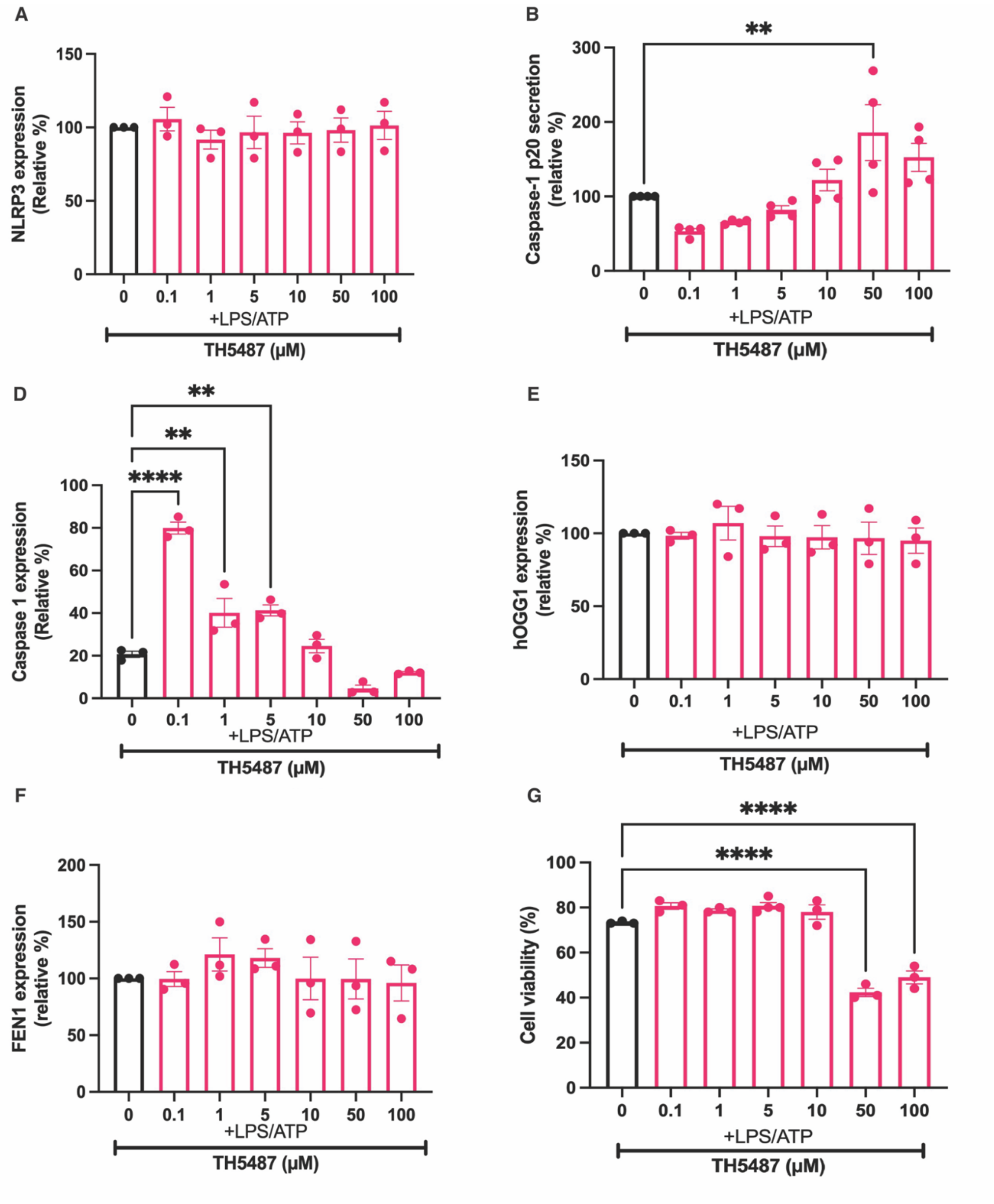

A
NLRP3 expression (Relative %)
150
100
50
0
0 0.1 1 5 10 50 100
+LPS/ATP
TH5487 (μM)
B
**
Caspase-1 p20 secretion (relative %)
300
200
100
0
0 0.1 1 5 10 50 100
+LPS/ATP
TH5487 (μM)
D
**
**
****
Caspase 1 expression (Relative %)
100
80
60
40
20
0
0 0.1 1 5 10 50 100
+LPS/ATP
TH5487 (μM)
E
hOGG1 expression (relative %)
150
100
50
0
0 0.1 1 5 10 50 100
+LPS/ATP
TH5487 (μM)
F
FEN1 expression (relative %)
200
150
100
50
0
0 0.1 1 5 10 50 100
+LPS/ATP
TH5487 (μM)
G
****
****
Cell viability (%)
100
80
60
40
20
0
0 0.1 1 5 10 50 100
+LPS/ATP
TH5487 (μM)

**figure S5: Other than caspase-1, treatment with TH5487 had no effect on other inflammasome-related proteins.** **(A)** Quantified relative amount of NLRP3 in the lysate of LPS/ATP stimulated immortalized mouse macrophages challenged with 0-100 µM TH5487. Error bars: mean +/- SEM, analyzed with one-way ANOVA. N=3 **(B)** Quantified relative amount of cxaspase-1 p20 in the supernatant of LPS/ATP stimulated immortalized mouse macrophages challenged with 0-100 µM TH5487. Error bars: mean +/- SEM, analyzed with one-way ANOVA. N=4, ** P = 0.0088 **(C)** Quantified relative amount of caspase-1 in the lysate of LPS/ATP stimulated immortalized mouse macrophages challenged with 0-100 µM TH5487. Error bars: mean +/- SEM, analyzed with one-way ANOVA. N=3, **** P<0.0001, **P= 0.0072 **(D)** Quantified relative amount of hOGG1 in the lysate of LPS/ATP stimulated immortalized mouse macrophages challenged with 0-100 µM TH5487. Error bars: mean +/- SEM, analyzed with one-way ANOVA. N=3 **(E)** Quantified relative amount of FEN1 in the lysate of LPS/ATP stimulated immortalized mouse macrophages challenged with 0-100 µM TH5487. Error bars: mean +/- SEM, analyzed with one-way ANOVA. N=3 **(F)** cell viability as measured by trypan blue of all cells treated and untreated. Error bars: mean +/- SEM, analyzed with one-way ANOVA N=3, **** P<0.0001

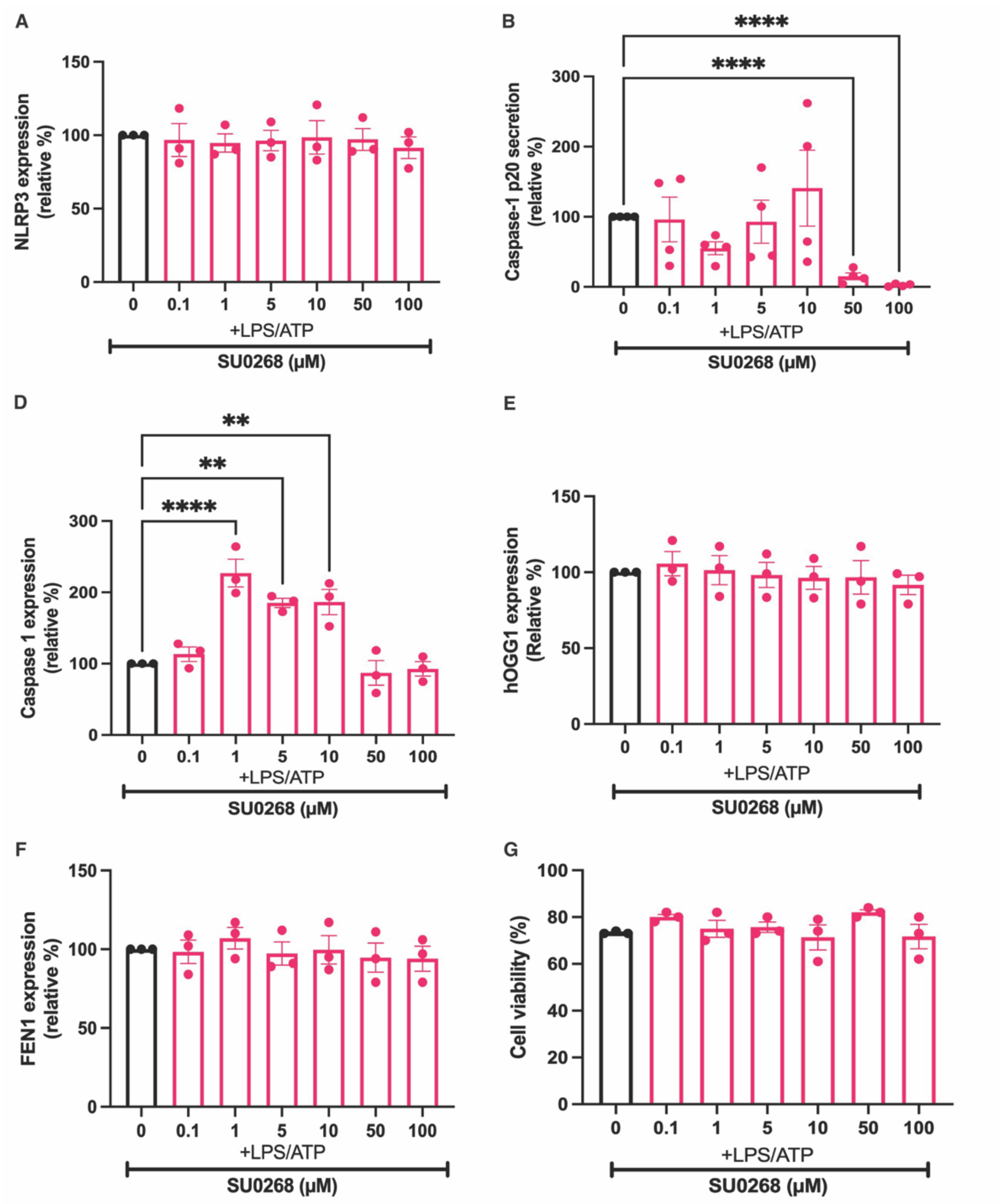
A
150
100
50
0
NLRP3 expression (relative %)
0
0.1
1
5
10
50
100
+LPS/ATP
SU0268 (μM)
B
****
****
300
200
100
0
Caspase-1 p20 secretion (relative %)
0
0.1
1
5
10
50
100
+LPS/ATP
SU0268 (μM)
D
**
**
****
300
200
100
0
Caspase 1 expression (relative %)
0
0.1
1
5
10
50
100
+LPS/ATP
SU0268 (μM)
E
150
100
50
0
hOGG1 expression (Relative %)
0
0.1
1
5
10
50
100
+LPS/ATP
SU0268 (μM)
F
150
100
50
0
FEN1 expression (relative %)
0
0.1
1
5
10
50
100
+LPS/ATP
SU0268 (μM)
G
100
80
60
40
20
0
Cell viability (%)
0
0.1
1
5
10
50
100
+LPS/ATP
SU0268 (μM)

**figure S6: Other than caspase-1, treatment with SU0268 had no effect on other inflammasome-related proteins. (A)** Quantified relative amount of NLRP3 in the lysate of LPS/ATP stimulated immortalized mouse macrophages challenged with 0-100μM SU0268. Error bars: mean +/- SEM, analyzed with one-way ANOVA. N=3 **(B)** Quantified relative amount of caspase-1 p20 in the supernatant of LPS/ATP stimulated immortalized mouse macrophages challenged with 0-100 μM SU0268. Error bars: mean +/- SEM, analyzed with one-way ANOVA. N=4, ****P = <0.0001 **(C)** Quantified relative amount of caspase-1 in the lysate of LPS/ATP stimulated immortalized mouse macrophages challenged with 0-100 μM SU0268. Error bars: mean +/- SEM, analyzed with one-way ANOVA. N=3, ****P <0.0001, **P = 0.0021 **(D)** Quantified relative amount of hOGG1 in the lysate of LPS/ATP stimulated immortalized mouse macrophages challenged with 0-100 μM SU0268. Error bars: mean +/- SEM, analyzed with one-way ANOVA. N=3 **(E)** Quantified relative amount of FEN1 in the

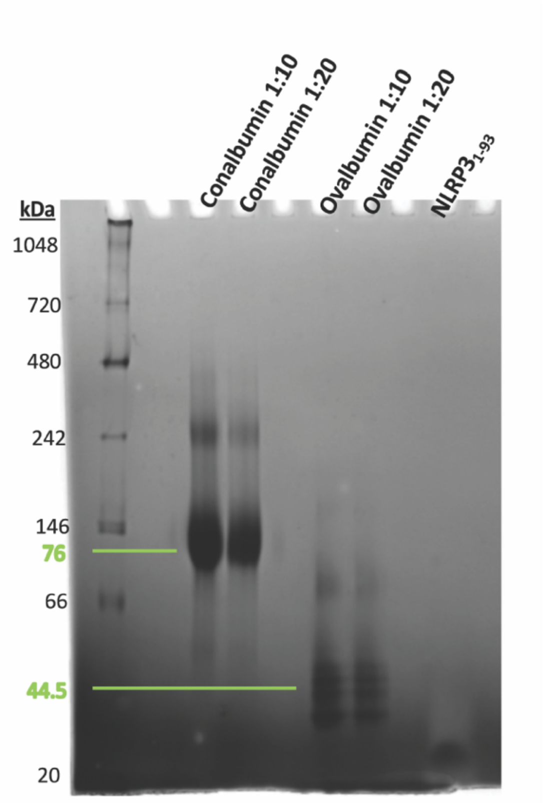


**figure S7. NLRP3$_{1\text{-}93}$ Purification.** Following affinity chromatography, size exclusion (SEC) was performed with a 16/600 Supersose 6 column. Expected Molecular weight (MW)14 kDa**.** 4-16% native Bis-Tris blue total protein gel on fraction E1 following SEC and dialysis. Conalbumin MW 76 kDa (top green line) and Ovalbumin MW 44.5 kDa (bottom green line).

| WT NLRP3: AA codes | hOGG1: AA codes | hOGG1 TH5487 (Distance (Å)) | hOGG1 ox-DNA (Distance (Å)) |
|---|---|---|---|
| Lys 2 | Gln/Lys 249 | 13.14 | 13.16 |
| Ala 4 | Ala 251 | 11.007 | 10.743 |
| Cys 8 | Cys 255 | 8.307 | 8.09 |
| Ala 11 | Ala 258 | 16.378 | 16.109 |
| Asp 16 | Asp 268 | 3.55 | 3.506 |
| Asp 19 | Asp 268 | 8.016 | 8.501 |
| Val 20 | Val 267 | 13.253 | 13.029 |
| Asp 21 | Asp 268 | 7.976 | 7.551 |
| His 28 | His 270 | 9.701 | 9.119 |
| Asp 31 | Asp 278 | 3.001 | 2.94 |
| Tyr 32 | Tyr 279 | 6.761 | 6.885 |
| Pro 34 | Pro 283 | 23.896 | 23.531 |
| Pro 42 | Pro 291 | 17.982 | 17.717 |
| Gln 45 | Gln 294 | 13.559 | 13.753 |
| Thr 46 | Thr 295 | 9.57 | 9.581 |
| Leu 54 | Leu 299 | 0.603 | 0.69 |
| Gly 63 | Gly 308 | 0.535 | 0.54 |
| Trp 68 | Trp 313 | 3.756 | 3.61 |
| Ala 69 | Ala 314 | 1.699 | 1.515 |
| Ala 71 | Ala 316 | 4.806 | 4.714 |
| Val 72 | Val 317 | 4.323 | 3.552 |
| Phe 75 | Phe 319 | 2.538 | 2.078 |
| Ala 77 | Ala 321 | 3.835 | 3.836 |
| Arg 80 | Arg 324 | 9.17 | 8.604 |

**table S1: Distances between alpha carbons of specified amino acids in NLRP3 compared to hOGG1.** NLRP3 pyrin$_{(1\text{-}91)}$ (PDBID: 7PZC) docked into hOGG1$_{(249\text{-}325)}$ with either Ox-DNA (PDBID: 1EBM) or TH5487 (PDBID: 6RLW). NLRP3 (PDBID 7PZC) with hOGG1 Ox-DNA (PDBID: 1EBM): RMSD between 16 pruned atom pairs is 1.132 angstroms; (across all 69 pairs: 11.975) NLRP3 with hOGG1 TH5487: RMSD between 17 pruned atom pairs is 1.147 angstroms; (across all 67 pairs: 10.790).

| Attribute | Score | Notes |
|---|---|---|
| **Models produced** | 1 | |
| **MolProbity** | 1.78 | Best: between 0 and 1 |
| **Sequence Identity** | 37.68% | |
| **Clash Score** | 7.68 | |
| **Ramachandran Favored** | 94.87% | |
| **Ramachandran Outliers** | 3.85% | C8 Cys, C54 Val, C34 Pro |
| **Rotamer Outliers** | 0% | |
| **Bad Bonds** | 0/626 | |
| **Bad Angles** | 21/844 | |
| **Cis Non-Proline** | Mar-75 | |
| **GMQE** | 0.29 | Worst=0, 1=best |
| **QMEANDisCo Global** | 0.33 +/- 0.1 | Worst=0, 1=best |

**table S2: Attributes of the SWISS-MODEL generated NLRP3 model based on hOGG1 bound to Ox-DNA**

| NLRP3 Model: AA codes | hOGG1: AA codes | hOGG1 TH5487 (Distance (Å)) | hOGG1 ox-DNA (Distance (Å)) |
|---|---|---|---|
| Lys 2 | Gln/Lys 249 | 0.155 | 0.124 |
| Ala 4 | Ala 251 | 0.478 | 0.09 |
| Cys 8 | Cys 255 | 0.368 | 0.342 |
| Ala 11 | Ala 258 | 0.29 | 0.05 |
| Asp 16 | Asp 268 | 17.575 | 17.212 |
| Asp 19 | Asp 268 | 13.018 | 12.796 |
| Val 20 | Val 267 | 6.215 | 6.139 |
| Asp 21 | Asp 268 | 6.772 | 6.344 |
| His 28 | His 270 | 5.655 | 5.279 |
| Asp 31 | Asp 278 | 4.428 | 4.138 |
| Tyr 32 | Tyr 279 | 4.123 | 4.022 |
| Pro 34 | Pro 283 | 2.755 | 3.202 |
| Pro 42 | Pro 291 | 10.158 | 10.513 |
| Gln 45 | Gln 294 | 0.615 | 0.119 |
| Thr 46 | Thr 295 | 0.234 | 0.128 |
| Leu 54 | Leu 299 | 0.103 | 0.029 |
| Gly 63 | Gly 308 | 0.043 | 0.11 |
| Trp 68 | Trp 313 | 0.068 | 0.102 |
| Ala 69 | Ala 314 | 0.047 | 0.074 |
| Ala 71 | Ala 316 | 0.432 | 0.116 |
| Val 72 | Val 317 | 0.591 | 0.544 |
| Phe 75 | Phe 319 | 0.676 | 0.196 |
| Ala 77 | Ala 321 | 0.336 | 0.282 |
| Arg 80 | Arg 324 | 2.127 | 0.064 |

**table S3: Distances between alpha carbons of specified amino acids in SWISS-MODEL projection of NLRP3 pyrin based on hOGG1 bound to ox-DNA docked into hOGG1.** hOGG1$_{(249-325)}$ with Ox-DNA (PDBID: 1EBM) or TH5487 (PDBID: 6RLW). NLRP3 model with hOGG1 TH5487: RMSD between 32 pruned atom pairs is 0.358 angstroms; (across all 70 pairs: 5.661). NLRP3 model with hOGG1 Ox-DNA: RMSD between 31 pruned atom pairs is 0.256 angstroms (across all 72 pairs: 5.884).

| Antibody | Source | Identifier |
|---|---|---|
| **NLRP3 (pyrin targeting)** | AdipoGen | Cat#: AG-20B-0014-C100 (Cryo-2) |
| **NLRP3 (NACHT targeting)** | Cell Signaling | Cat#: 15101 (D4D8T) |
| **Cleaved-IL-1β (Asp117)** | Cell Signaling | Cat#: 63124 (E7V2A) |
| **Caspase-1 (p20)** | AdipoGen | Cat#: AG-20B-0042-C100 (Casper-1) |
| **OGG1** | Santa Cruz Biotechnology | Cat#: sc-376935 (G-5) |
| **FEN1** | Santa Cruz Biotechnology | Cat#: sc-28355 (B-4) |
| **beta-Actin** | Santa Cruz Biotechnology | Cat#: sc-47778 (C4) |
| **Anti-rabbit IgG, HPR-linked** | Cell Signaling | Cat#: 7074 |
| **Anti-mouse IgG, HPR-linked** | Cell Signaling | Cat#: 7076 |
| **Streptavidin, HRP-linekd** | BD Pharmingen | Cat#: 554066 |

**Table S4: Antibodies during for these studies.**
